# Novel 5-Hydroxymethylcytosine Markers for Pancreatic Cancer

**DOI:** 10.1101/425983

**Authors:** Chang Zeng, Zhou Zhang, Jun Wang, Brian C-H. Chiu, Lifang Hou, Wei Zhang

## Abstract

**OBJECTIVES:** Robust biomarkers for pancreatic cancer (PaC) early detection/prognosis are critical for improved patient survival. Our goal was to explore the biomarker potential of 5-hydroxymethylcytosines (5hmC), an epigenetic marker with a distinct role in cancer pathobiology, yet under-investigated due largely to technical constraints.

**METHODS:** We used the TAB-Array assay, a state-of-the-art technology to directly profile 5hmC at single base resolution with the Illumina EPIC array (>850,000 CpG sites) in 17 pairs of tumor/adjacent tissue samples from US patients.

**RESULTS:** We demonstrated distinctive distributions of 5hmC in tissues, and substantial differences between tumor and adjacent tissues, suggesting their diagnostic/prognostic value of for PaC.

**CONCLUSION:** This study established the potential of 5hmC as a novel epigenetic biomarker for PaC.

## 1. Background

Pancreatic cancer (PaC) ranks the fourth in the United States and eighth worldwide for cancer mortality, in part because it is usually detected when it is no longer surgically resectable [21]. While, early detection and prevention is the only viable option to improve survival that is critical for increased chances of survival but is currently unavailable for PaC [4]. Due to the low incidence of PaC in the population, most positive screening tests are likely false positives. Therefore, it remains a grand challenge to identify more effective biomarkers for early diagnosis, prognosis prediction, and precise management of this fatal disease.

Though the rapid advancement of high-throughput sequencing technologies has allowed the characterization of tumor related mutations, the low mutation frequency and lack of information on tissue of origin hampers detection sensitivity and specificity based on mutations alone. Recent studies demonstrated aberrant cytosine modifications in tumors, suggesting the potential applications of these epigenetic markers in cancer early detection and diagnosis [12, 15]. In particular, 5-hydroxymethylcytosine (5hmC), which is generated by oxidation of the more common 5-methylcytosines (5mC) through the Tet (ten-eleven translocation) family of enzymes, have been demonstrated as a stable modified cytosine in the human genome, covering ~0.1-1% of all CpG dinucleotides, with a distinct gene regulatory role and genomic distribution from 5mC [22, 23, 27]. 5hmC has been implicated in various human cancers, and the global reduction of 5hmC has been observed in both solid tumors and hematological malignancies [13, 20, 24]. Therefore, given their prevalence in the human genome, gene regulation relevance, and biochemical stability, the 5hmC markers have the potential as a novel class of cancer biomarkers for diagnosis and prognosis [14].

Despite the remarkable potential of 5hmC as cancer biomarkers, their uses in clinical samples have been limited due mainly to the lack of enabling technologies that can distinguish 5hmC from other modified cytosines. For example, although the Illumina Infinium arrays have become a widely used epigenomic platform, the conventional bisulfite-conversion protocol cannot distinguish 5hmC from 5mC. In the current study, therefore, we used the TAB-Array, a state-ofthe-art technique that is implemented on the Illumina EPIC array (covering ~850,000 CpG modification sites), to map and profile 5hmC levels directly in paired tumor and adjacent tissues collected from 17 US patients with primary pancreatic adenocarcinoma [19, 26]. We demonstrated the potential of targeting 5hmC as a novel epigenetic biomarker for PaC, thus laying the foundation for future development of clinically convenient assays for early detection of PaC, for example using emerging technologies for liquid biopsies [11].

## 2. Materials and Methods

### 2.1 Study Subjects

De-identified clinically annotated leftover tissues from 17 adult patients with primary pancreatic adenocarcinoma (females: n = 10, males: n = 7; white: n = 7, black: n = 1, hispanic: n = 1, other: n = 8; age at diagnosis [mean, standard deviation]: 62.5±15.8 yrs) were obtained from the Human Tissue Resource Center of The University of Chicago Medical Center. Patients treated with chemotherapy, radiation therapy or immunotherapy were excluded from this pilot study. Diagnosis of primary pancreatic adenocarcinoma was histologically confirmed. The Institutional Review Board at each institute approved this study.

### 2.2 DNA Isolation and TAB-Array Profiling

Genomic DNA (gDNA) was then isolated using the DNeasy Blood & Tissue kit (Qiagen, Germany), purified, and quantified, followed by the TAB-Array assay at the University of Chicago Genomics Core Facility [12]. Briefly, gDNA prepared from each sample was split into two fractions. One fraction was bisulfite converted and analyzed by hybridization to the Illumina EPIC array (covering > 850,000 CpG modification sites that encompass the entire human genome). The second fraction was first glucosylated to protect the 5hmC from oxidation by recombinant Tet1 enzyme using the TAB-Array kit (WiseGene, Inc., Chicago, IL), followed by bisulfite treatment and hybridization to the EPIC array. Unmethylated cytosines can be converted to thymidines by bisulfite treatment and amplification. In the absence of TAB treatment, both 5hmC and 5mC are protected and remain as cytosines. TAB treatment thus protects 5hmC, while 5mC is oxidized and converted by sodium bisulfite to thymidines. The 5hmC sites was distinguished from 5mC by comparing the hybridization results in the two fractions for each tissue sample. The raw and processed array data have been deposited into the NCBI/Gene Expression Omnibus database.

### 2.3 Processing of the TAB-Array Data and Detection of Differential Loci

The raw TAB-Array and conventional EPIC array data were processed using ENmix R package, separately [15]. Briefly, After QC process, 3,266 (TAB-Array) and 25,088 (conventional bisulfite conversion) low quality CpGs (detection p-value > 10-6 in more than 5% samples) were detected for each assay, respectively. The detected low quality CpGs were pooled and removed from both datasets. For each dataset, the modification signal intensities were modeled with a flexible exponential-normal mixture distribution and the background noise was modeled with a truncated normal distribution for background correction. Quantile normalization of modification intensities was performed across all 34 samples, and the β-values for each CpG site were then obtained. The final 5hmC and 5mC β-values were estimated using the Maximum Likelihood Estimate (MLE) from paired bisulfite conversion and TAB-treated samples. The CpGs on sex chromosome were removed, resulting 822,760 CpGs in the final datasets for further statistical analysis.

To detect differentially modified 5hmc loci across the 17 paired tumor and adjacent tissue samples, the R package Limma was used to perform a paired t-test for each probe [16]. A Bonferroni-corrected p-value less than 0.05 was considered significant.

### 2.4 Evaluating Genomic Features and Functional Relevance

To characterize the distribution of tumor-associated 5hmC loci, the genomic coordinates of these 5hmC loci were extracted and mapped to their respective host genes based on the current GENCODE annotations (hg19) [7]. We also mapped the tumor-associated 5hmC loci to known histone modification marks derived from pancreas using ChIP-seq in the Roadmap Epigenomics Project [1]. To measure the enrichment of these tumor-associated 5hmC loci with various genomic features (e,g., histone modification marks), the one-sided Fisher’ exact test was performed to calculate the odds ratio and p-value using the EPIC array as reference. The biological relevance of these 5hmC-containing genes was explored using the NIH/DAVID tool for any enriched Kyoto Encyclopedia of Genes and Genomes (KEGG) pathways and Gene Ontology (GO) biological processes [5, 9, 10].

### 2.5 Exploring Prognostic Value of 5hmC for PaC

The Human Pathology Atlas (HPA) is a database that explores the prognostic role of protein-coding genes in 17 different cancers by transcriptomics and antibody-based profiling [25]. The HPA predicts prognostic role of a particular gene for PaC using The Cancer Genome Atlas (TCGA) project where transcriptomics data are available from 176 US patients with pancreatic adenocarcinoma [25]. According to the HPA, the “favorable” (or “unfavorable”) gene was defined as those genes with higher relative expression levels at diagnosis and in the meantime corresponded to higher (or lower) five-year overall survival, and the log-rank p-value based on five-year survival was used for statistical significance to infer potential prognostic value of the 5hmC-contatining marker genes detected in the tumors. To determine whether the PaC-associated 5hmC were enriched in genes with prognostic value, 5hmC loci were randomly sampled from the EPIC array background for 100,000 times, and then the number of genes with prognostic value based on the HPA was counted.

## 3. Results

### 3.1 Genomic Distribution of 5hmC in PaC Tissues

Using the TAB-Array assay, we profiled 5hmC levels in 17 pairs of tumor and adjacent tissue samples. Consistent with previous epigenetic studies using the Infinium arrays [17], the detected 5mC in PaC tissues showed a two-mode distribution with peaks at both lower and upper end of modification levels (<0.20 and >0.80 in terms of β-value). In contrast, the 5hmC modification sites showed a unimodal distribution enriched at the lower end of modification levels (**Fig.1A**). Notably, the 5hmC and 5mC levels were inversely correlated for the majority of the CpGs, while only ~ 20% of the CpGs demonstrated weak positive correlations (**Fig.1B**), indicating dramatic epigenetic changes associated with tissue pathology. Further separation of the modification sites into various genomic features showed the enrichment of high 5mC modification levels within CpG islands (**Fig.1C**) but in gene bodies for 5hmC, indicating distinct genomic distributions for these two types of modified cytosines (**Fig.1D**).

**Fig.1.**
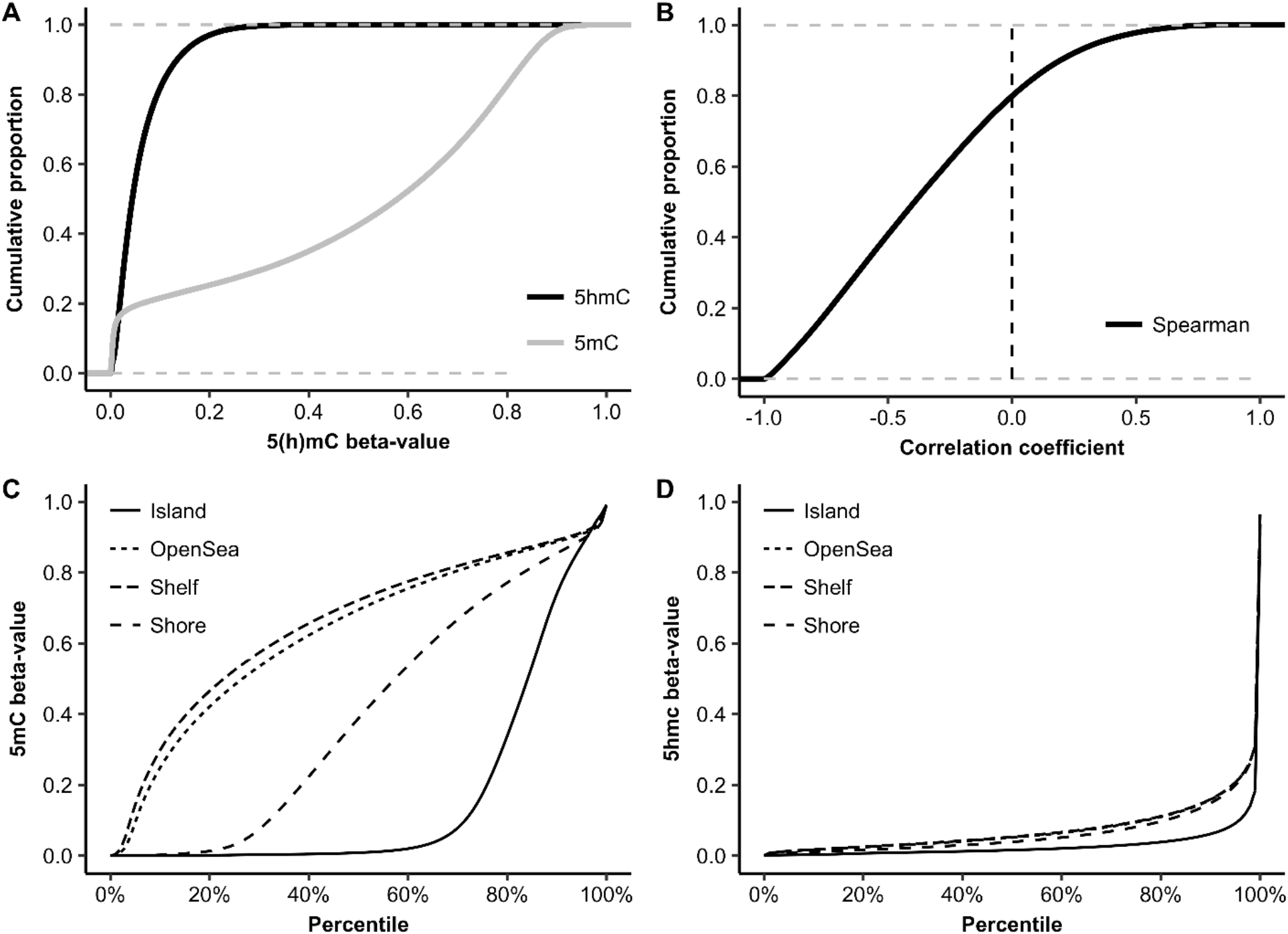
The Profiled 5hmC Loci Show a Distinct Genomic Pattern. In contrast to 5mC, in general, 5hmC loci are biased toward lower end of modification levels, and show a distinct genomic distribution pattern, enriched in gene bodies. (A) Comparison of the cumulative proportions between 5hmC and 5mC in terms of modification level (β-value); (B) 5hmC modifications are in general, negatively correlated with 5mC; Comparison of the genomic distributions between (C) 5mC and (D) 5hmC in PaC tissues.

### 3.2 PaC-Associated 5hmC Loci

We identified 1,118 differential 5hmC loci among the 17 pairs of tumor and adjacent tissue samples with the Bonferroni-corrected p-value < 0.05 (**Fig.2A**). Examining the genomic distribution pattern of these PaC-associated 5hmC loci suggested an over-representation in the genic region, particularly the gene bodies (**Fig.2B**). Enrichment analysis showed that the differential 5hmC CpGs were more likely to reside in intronic regions while less likely to reside in promoter regions and exons. Based on the Roadmap Epigenomics Project data derived from pancreas, we further demonstrated that our detected PaC-associated 5hmC loci were enriched with histone modification marks for enhancers and active gene expression regulation (**Fig.2B**). Specifically, we observed significant enrichment in enhancers (marked by H3K4me1), regions with active transcription (marked by H3K27ac), gene bodies (marked by H3K36me3), but depletion in promoters (marked by H3K4me3), regions with polycomb repression (H3K27me3), and formation of heterochromatin (H3K9me3) (**Fig.2B**).

**Fig.2.**
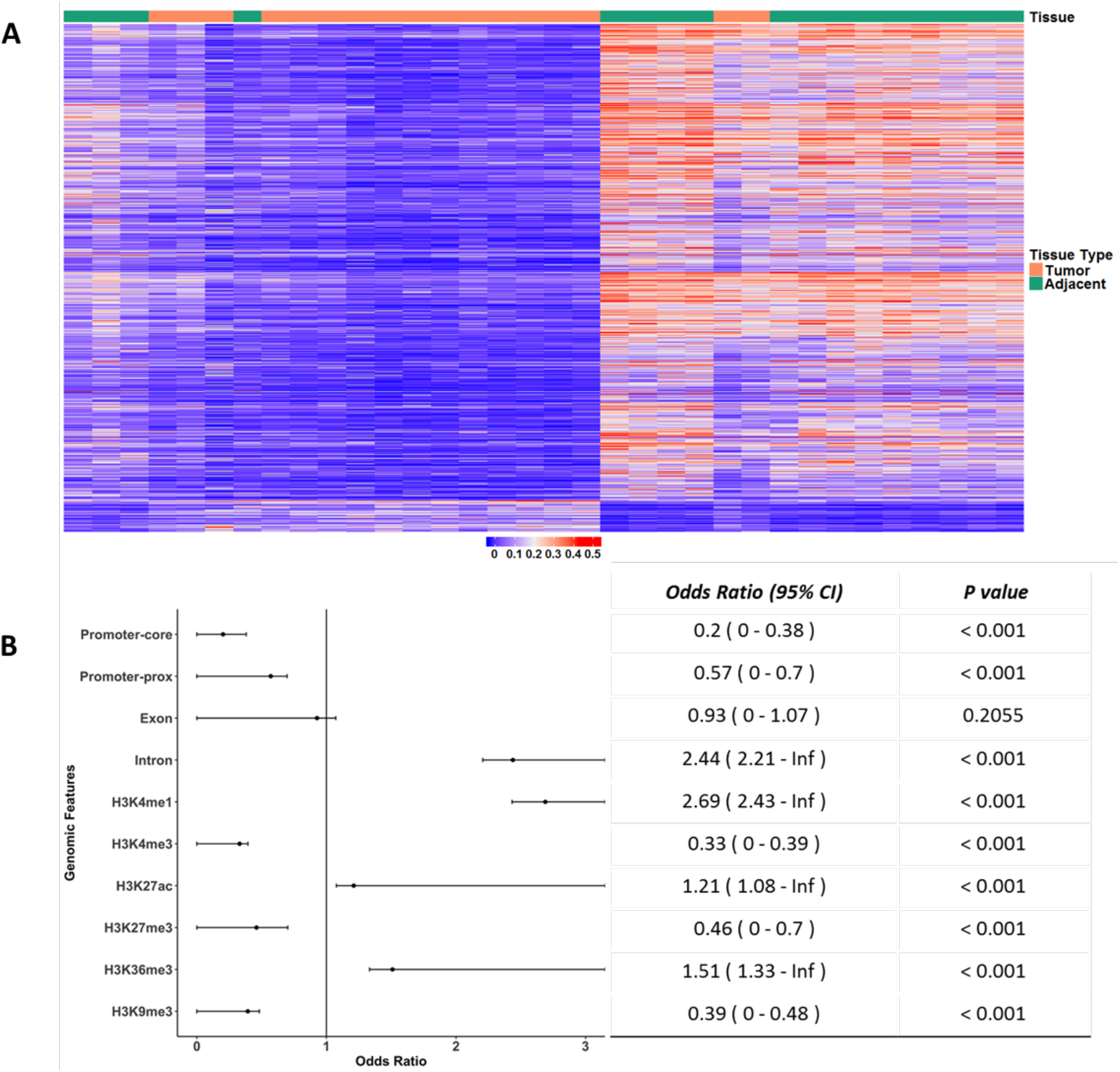
PaC-Associated 5hmC Loci. (A) The heatmap is plotted using the top 1,118 differential 5hmC loci (Bonferroni-corrected p-value < 0.05) (Supplemental Table 1); (B) The detected PaC-associated 5hmC loci are enriched with gene bodies and histone modification marks for enhancers and active gene expression derived from pancreas in the Roadmap Epigenomics Project.

### 3.3 Exploring Functional Relevance and Prognostic Value

We searched the KEGG database and GO to explore functional relevance of the detected PaC-associated 5hmC loci using the NIH/DAVID tool. We found that genes containing the differential 5hmC loci are enriched in canonical pathways relevant to cancer pathobiology, such as PI3KAkt signaling pathway, Rap1 signaling pathway, and Ras signaling pathway; as well as GO biological processes, such as negative regulation of Wnt receptor signaling pathway and lipid metabolic process (**Fig.3A**) [2, 16, 18]. We further explored potential prognostic value of those genes containing differential 5hmC sites between PaC tumor and adjacent tissues, based on the five-year survival data on PaC in the HPA database. The permutation test on 822760 CpGs sites with the EPIC array as background for 100,000 times suggested that down-regulated 5hmC loci were enriched in genes with unfavorable prognostic values in PaC (p-value = 0.0123) (**Fig.3B**), indicating their potential for prognostic prediction as well.

**Fig.3.**
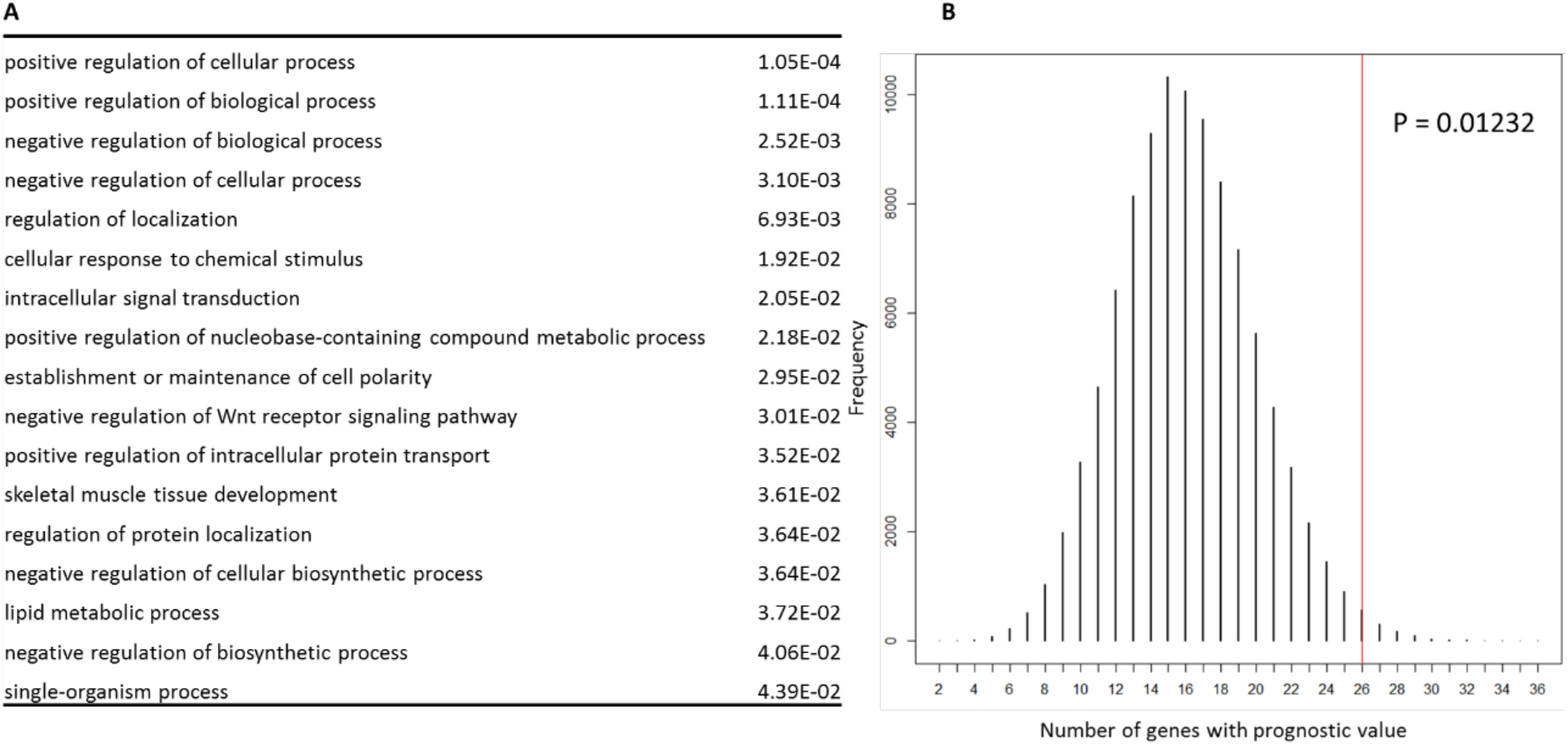
Functional Relevance and Prognostic Value of PaC-Associated 5hmC Loci. (A) The host genes containing PaC-associated 5hmC loci are enriched with certain KEGG pathways and GO biological processes relevant to cancer pathobiology; (B) Evaluation of prognostic value of PaC-associated 5hmC loci using a permutation test, based on the Human Pathology Atlas database. The real data point is shown as a red vertical line.

## 4. Discussion

Despite great efforts to improve the clinical outcomes of PaC, PaC is expected to become the second leading cause of cancer death in the United States by 2030[21]. Therefore, effective biomarkers are urgently needed for PaC diagnosis, prognosis, and disease surveillance to improve survival. Epigenetic regulators are particularly promising biomarkers for early detection of PaC, considering their tissue specificity, quantitative nature, availability of high-throughput technologies. In this pilot study, using the TAB-Array, a recently developed technique, we profiled directly 5hmC, the currently under-studied cytosine modification but with demonstrated cancer relevance, in paired tumor and adjacent tissue samples from patients with PaC. Our data showed a distinct genome-wide distribution pattern and modification landscape for 5hmC (**Fig.1**), consistent with previous studies [3]. Notably, we found that the detected 5hmC are in general inversely correlated with 5mC modifications, consistent with their putatively different gene regulatory roles, and supporting the importance of distinguishing 5hmC from 5mC in epigenetic biomarker discovery.

Specifically, by comparing the 5hmC profiles between tumor and adjacent tissue samples, we identified a substantial proportion of 5hmC loci that are differentially modified (**Fig.2**). Consistent with their putative gene regulatory roles, these differential 5hmC CpG sites were significantly over-represented in pancreas-derived histone modification marks for enhancers and active gene expression. Moreover, cancer related pathways such as Ras signaling pathway, PI3K-Akt signaling pathway were enriched in the PaC-associated 5hmC sites. Further examination of the HPA database suggested an over-representation of genes with prognostic value for five-year survival among our PaC-associated genes (**Fig.3B**), including for example genes such as *HDAC4*, *Rab1, FIP3*, which are known to be related to PaC pathobiology and/or metastasis[6, 8]. Taken together, our findings from this pilot study indicated potential biological and functional relevance of those genes containing the detected differential 5hmC in PaC tissues.

## 5. Conclusion

In this pilot study of 17 pairs of PaC tumor and adjacent tissue samples, we established the feasibility of using TAB-Array to profile 5hmC in PaC tissues. A substantial proportion of 5hmC loci were found to be differentially modified between tumor and adjacent tissues from PaC patients. Finally, though limited by sample size and clinical data, our pilot study showed functional relevance of those genes containing PaC-associated 5hmC and suggested potential prognostic value of 5hmC markers. Overall, based on our findings, 5hmC is warranted to be an important epigenetic target in future biomarker discovery studies for PaC to improve diagnosis and prognosis.

